# Development of A Novel Highly Selective TLR8 Agonist for Cancer Immunotherapy

**DOI:** 10.1101/2020.03.14.991760

**Authors:** Yuxun Wang, Heping Yang, Huanping Li, Shuda Zhao, Yikun Zeng, Panpan Zhang, Xiaoqin Lin, Xiaoxiang Sun, Longsheng Wang, Guangliang Fu, Yaqiao Gao, Pei Wang, Daxin Gao

## Abstract

Toll-like receptors (TLRs) are a family of proteins that recognize pathogen associated molecular patterns (PAMPs). Their primary function is to activate innate immune responses while also involved in facilitating adaptive immune responses. Different TLRs exert distinct functions by activating varied immune cascades. Several TLRs are being pursued as cancer drug targets. We discovered a novel, highly potent and selective small molecule TLR8 agonist DN052. DN052 exhibited strong in vitro cellular activity with EC50 at 6.7 nM and was highly selective for TLR8 over other TLRs including TLR4, 7 and 9. The selectivity profile distinguished DN052 from all other TLR agonists currently in clinical development. DN052 displayed excellent in vitro ADMET and in vivo PK profiles. DN052 potently inhibited tumor growth as a single agent. Moreover, combination of DN052 with the immune checkpoint inhibitor, selected targeted therapeutics or chemotherapeutic drugs further enhanced efficacy of single agents. Mechanistically, treatment with DN052 resulted in strong induction of pro-inflammatory cytokines in ex vivo human PBMC assay and in vivo monkey study. GLP toxicity studies in rats and monkeys demonstrated favorable safety profile. This led to the advancement of DN052 into phase I clinical trials.

## INTRODUCTION

Human immune defense system comprises both innate and adaptive immune pathways (1). Most of the targets drugged by the recently approved cancer immunotherapeutic agents including the immune checkpoint proteins PD-1, PD-L1 and CTLA-4 function in adaptive immune pathways (2, 3). In contrast, targets involved in the innate immune pathway had been under-developed (4, 5). Innate immunity acts as the body’s first line of immune defense. Drugs targeting innate immunity hold potential for more rapid and broader spectrum anti-cancer effect than adaptive immunity. Furthermore, combinations of drugs targeting innate and adaptive immunity are expected to produce strong synergistic efficacy owing to their complementary nature as body’s immune defense (6). Toll-like receptors (TLRs) are a family of proteins that recognize pathogen associated molecular patterns (PAMPs). Their primary function is to activate innate immune responses while they are also involved in facilitating adaptive immune responses (7). Different TLRs differ in their expression in various target cells and exert distinct functions by activating varied immune cascades (7-9). In the TLR family, several TLRs have been studied as cancer drug targets such as TLR2, 4, 7, 8 and 9 (7, 10-15). Some of them are being drugged by conventional small molecule modality while others are being targeted by unconventional molecules such as oligonucleotide agonists for TLR9 (13, 14). We reasoned that among different TLRs amenable to small molecule modality, specific targeting TLR8 would be advantageous. First, from the efficacy perspective, TLR8 has been shown to be necessary and sufficient to reverse the immune suppressive function of Treg cells leading to strong tumor inhibition (16-20). TLR8 activation has also been shown to induce apoptosis of MDSCs (21). MDSCs are another major type of cells that suppress immune response. Apoptosis of MDSCs can lead to activated and enhanced immune response to tumors. Treg and MDSC are the major cause of failure in cancer immunotherapy (21-23). Therefore, reversing Treg and MDSC mediated immune suppression by activating TLR8 can elicit potent immune response resulting in strong anti-cancer effect (16, 24). Moreover, TLR8 has been shown to induce terminal differentiation-mediated tumor suppression in acute myeloid leukemia (AML) (25). Second, from safety perspective, TLR8 agonists have better safety profile allowing systemic administration whereas targeting other TLRs such as TLR4, 7, TLR7/8 dual, TLR9 appear to be less tolerable when dosed systemically and are mainly dosed locally through intra-tumoral injection or limited to topical use (10, 13). For example, imiquimod, also known as Aldara, an approved drug predominantly targeting TLR7 is used as a topical drug because it is too toxic to use systemically, which dramatically limits its clinical application (26-28). Systemic administration of TLR7 agonists has been explored using relatively low doses or prodrug form to reduce their systemic toxicity. However, further optimization of their dosing regimen and the compounds is necessary before systemic dosing can be applied in the clinic (29-31). TLR7/8 dual agonists are being developed and also administered by intra-tumoral injection (32, 33). Motolimod is generally known as a TLR8 agonist (34). However, consistent with literature reports, we have found motolimod has weak activity on TLR7 in addition to its agonistic activity over TLR8 indicating it is in fact a TLR7/8 dual agonist though it predominantly targets TLR8 (35). Motolimod is administered subcutaneously, a systemic dosing route (36, 37). However, the clinical doses used for motolimod were relatively low and its efficacy appeared to be limited suggesting motolimod had a narrow therapeutic index (36, 37). Nevertheless, unlike other TLR7/8 dual agonists, motolimod can be dosed systemically suggesting its weaker activity on TLR7 relative to other TLR7/8 dual agonists might contribute to its improved tolerability. TLR9 agonists are oligonucleotide based molecules that are distinct from conventional small molecules and are also dosed by intra-tumoral injection (14). Taken together, among different TLRs, specific targeting TLR8 is expected to be advantageous considering both efficacy and safety characteristics.

However, there are currently no approved drugs for TLR8 selective agonists. As described above, motolimod, a known TLR8 agonist, is in fact a TLR7/8 dual agonist albeit its TLR7 activity is much weaker than its activity on TLR8 based on in vitro cell-based assays (35). Motolimod is currently in phase 1/2 clinical development for cancer indications (34, 36-41). Another known TLR8 agonist in clinical development is selgantolimod (GS-9688). Interestingly, GS-9688 also has activity over TLR7 in addition to its predominant activity on TLR8 (Daffis, et al. EASL 2017 Amsterdam Netherlands). GS-9688 is now in Ph 2 development for chronic hepatitis B (hepatitis B infection) indication. GS-9688 is administered orally. However, the bioavailability of GS-9688 is extremely poor restricting its exposure to the gut, which is purposely designed to avoid its severe toxicity caused by systemic exposure (Daffis, et al. EASL 2017 Amsterdam Netherlands).

Therefore, there is an urgent need for a truly selective TLR8 agonist that would distinguish itself from all other TLR agonists. Through structure-based drug design, we discovered a novel, highly potent and selective small molecule TLR8 agonist with no activity on TLR7, namely DN052 for cancer indications. Since motolimod is a closely related TLR8 agonist in clinical development for cancer indications we included motolimod as a reference benchmark molecule in our study. Here, we present strong evidence demonstrating DN052 is a novel TLR8 selective agonist differentiated from other known TLR8 agonists and DN052 possesses excellent drug-like properties and is advancing in phase 1 clinical development.

## MATERIALS AND METHODS

### In vitro experiments

HEK-Blue™ hTLR4, 7, 8 and 9 cell lines were purchased from InvivoGen (Hong Kong). The cells express the human TLR gene and NF-κB/AP-1-inducible SEAP (secreted embryonic alkaline phosphatase) reporter gene. SEAP levels produced upon TLR stimulation can be determined by QUANTI-Blue™. In the cell-based assay, the main reagents used were QUANTI-Blue™ (InvivoGen) and ATPlite 1 Step (Perkin Elmer). The main instruments used were the microplate reader SpectraMax 340PC (Molecular Device) and Envision (Perkin Elmer). Motolimod and DN052 were synthesized in house. The compounds were in 10 mM stock solution in DMSO and stored at −20°C. The compounds were serially diluted in the concentration range from 0.5 nM to 15 µM (up to 50 μM in the hTLR4, hTLR7 and hTLR9 assays) and added to 96-well plates. DMSO was used as the negative control. The cells were cultured and treated with the compounds at 37°C, 5% CO_2_ for 24 hours. After 24 hours of incubation, 20 μl of the supernatant of each well was added to 180 μl of QUANTI-Blue™ (pre-warmed at 37°C). The plates were incubated at 37°C for 1.5 hour. After 1.5 hours of incubation, the optical density was measured using the spectrophotometer at 650 nm in the microplate reader of SpectraMax 340PC. The cell viability of each well was determined using ATPlite 1 Step following the manufacturer’s instruction. The luminescence signal in each well of plates was measured in the microplate reader of Envision.

For the off-target screen, 10 µM of compounds were tested in Eurofins Cerep44 panel following the manufacturer’s instruction. Compound binding was calculated as a % inhibition of the binding of a radioactively labeled ligand specific for each target. Compound enzyme inhibition effect was calculated as a % inhibition of control enzyme activity.

### DMPK and hERG assays

CYP inhibition assay was conducted to evaluate the inhibitive potential of the compounds on CYP 1A2, 2C9, 2C19, 2D6 and 3A4 using human liver microsomes (BD Gentest). Curve-fitting was performed to calculate IC50 using a Sigmoidal (non-linear) dose-response model using GraphPad Prism (GraphPad Software Inc.).

To determine the in vivo pharmacokinetic profiles, the compounds were formulated with 11% captisol and then administered iv, ip, sc and po into Sprague Dawley rats or iv and sc into mice or cynomolgus monkeys, respectively. Blood samples were collected before dosing and 0.083 h,0.25 h,0.5 h,1 h,2 h,4 h,6 h,8 h,12 h and 24 h after dosing. Blood samples were collected in tubes containing K_2_EDTA. Plasma was isolated from the blood samples by centrifugation of the blood samples at 2400 x g for 5 min at 4 °C. Plasma samples were measured using LC-MS/MS (Waters ACQUITY UPLC and TQ 6500+) to determine the concentration of compounds. The data was analyzed using WinNonlin 7.0 to calculate the PK parameters such as AUC0-t, AUC0-∞, MRT0-∞, Cmax, Tmax, t_1/2_ and F.

hERG assay was conducted using CHO cell line stably transduced with hERG cDNA. Whole-cell recordings were performed using automated QPatch (Sophion Biosciences). Data were analyzed using Assay Software provided by Sophion (assay software V5.0), Microsoft Excel and Graphpad Prism.

### In vivo efficacy studies

All the animal experiments were conducted following Association for Assessment and Accreditation of Laboratory Animal Care (AAALAC)’s guideline. The animal protocols were approved by the Institutional Animal Care and Use Committee (IACUC). Cell lines were purchased from credible vendors including ATCC. After tumor cell inoculation, the animals were checked daily for morbidity and mortality. At the time of routine monitoring, the animals were checked for any effects of tumor growth and treatment on the animals’ well-being such as mobility, food and water consumption, body weight gain/loss (body weights were measured twice a week), matting and any other abnormalities. Clinical observations were recorded. Tumor volumes were measured twice a week in two dimensions using a caliper and the volume was expressed in mm^3^ using the formula: V = 0.5 a × b^2^ where a and b are the long and short diameters of the tumor, respectively. The procedures of dosing as well as tumor and body weight measurement were conducted in a laminar flow cabinet. For CT26 experiments, CT26 mouse colon cancer cells were cultured in vitro as a monolayer culture in RPMI1640 medium supplemented with 10% fetal bovine serum and 1% penicillin/streptomycin at 37°C in an atmosphere of 5% CO2 in air. The CT26 cells were routinely passaged twice per week by trypsin-EDTA treatment. The cells growing in the exponential growth phase were harvested and counted for tumor inoculation. 6-8 weeks old female Balb/c mice were purchased from Shanghai Lingchang Bio-Technology Co. Ltd. (Shanghai, China). Each mouse was inoculated subcutaneously at the right flank region with 1×10^5^ CT26 tumor cells in 100 µl ice-cold PBS. The tumor-bearing mice were randomized into different groups with 6 mice per group. Treatment started when the mean tumor volume reached 62 mm^3^. DN052 was formulated in 11% captisol and then subcutaneously (sc) administered once a week to the tumor-bearing mice at 40, 80 and 160 mg/kg, respectively. In the combination study in CT26 model, the mice were randomized into groups with 7 mice per group. Treatment started when the mean tumor volume reached 54 mm^3^. DN052 was sc administered at 40 mg/kg once a week. 50 mg/kg of cyclophosphamide (CTX) was intraperitoneally (ip) dosed once a week to CT26 tumor-bearing mice. Vehicle treated mice were included as controls.

For EMT6 experiments, EMT6 mouse breast cancer cells were cultured in vitro as a monolayer culture in RPMI1640 medium supplemented with 10% fetal bovine serum and 1% penicillin/streptomycin at 37°C in an atmosphere of 5% CO2 in air. The tumor cells were routinely passaged twice per week by trypsin-EDTA treatment. The cells growing in the exponential growth phase were harvested and counted for tumor inoculation. 6-8 weeks old female Balb/c mice were purchased from Beijing Vital River Laboratory Animal Technology Co., Ltd. (Beijing, China). Each mouse was inoculated subcutaneously at the right flank region with 1×10^5^ EMT6 tumor cells in 100 µl ice-cold PBS. The tumor-bearing mice were randomized into different groups with 8 mice per group. Treatment started when the mean tumor volume reached 58 mm^3^. DN052 was sc administered once a week to the tumor-bearing mice at 40, 80 and 160 mg/kg, respectively. Motolimod was sc administered once a week at 40 mg/kg. In the combination study in EMT6 model, the mice were randomized into groups with 6 mice per group. Treatment started when the mean tumor volume reached 58 mm^3^. DN052 was sc administered at 40 mg/kg once a week. AZD-1775 was dosed at 30 mg/kg, oral (po), bid to EMT6 tumor-bearing mice. Vehicle treated mice were included as controls.

For H22 experiment, H22 mouse liver cancer cells were cultured in vitro as a monolayer culture in RPMI1640 medium supplemented with 10% fetal bovine serum and 1% penicillin/streptomycin at 37°C in an atmosphere of 5% CO2 in air. The tumor cells were routinely passaged twice per week by trypsin-EDTA treatment. The cells growing in the exponential growth phase were harvested and counted for tumor inoculation. 6-8 weeks old female Balb/c mice were purchased from Beijing Vital River Laboratory Animal Technology Co., Ltd. (Beijing, China). Each mouse was inoculated subcutaneously at the right flank region with 6×10^5^ H22 tumor cells in 100 µl ice-cold PBS. The tumor-bearing mice were randomized into different groups with 8 mice per group. Treatment started when the mean tumor volume reached 72 mm^3^. DN052 was sc administered at 40 mg/kg once a week. Sorafenib was administered at 10 mg/kg, po, bid. AZD-1775 was dosed at 30 mg/kg, po, bid. Vehicle treated mice were included as controls.

For MC38 experiment, MC38 mouse colon cancer cells were cultured in vitro as a monolayer culture in DMEM medium supplemented with 10% fetal bovine serum and 1% penicillin/streptomycin at 37°C in an atmosphere of 5% CO2 in air. The tumor cells were routinely passaged twice per week by trypsin-EDTA treatment. The cells growing in the exponential growth phase were harvested and counted for tumor inoculation. 7-8 weeks old female C57BL/6 mice were purchased from Beijing Vital River Laboratory Animal Technology Co., Ltd. (Beijing, China). Each mouse was inoculated subcutaneously at the right flank region with 5×10^5^ MC38 tumor cells in 100 µl ice-cold PBS. The tumor-bearing mice were randomized into different groups with 7 mice per group. Treatment started when the mean tumor volume reached 57 mm^3^. DN052 was sc administered at 40 mg/kg once a week. The anti-PD-1 monoclonal antibody αPD-1 (Clone RMP1-14, Bioxcell) was administered at 10 mg/kg, ip, once a week. Vehicle treated mice were included as controls.

For HL-60 experiment, HL-60 human Acute Promyelocytic Leukemia (AML) cells were cultured in vitro in suspension in RPMI1640 medium supplemented with 10% fetal bovine serum and 1% penicillin/streptomycin at 37°C in an atmosphere of 5% CO2 in air. The tumor cells were routinely passaged twice per week. The cells growing in the exponential growth phase were harvested and counted for tumor inoculation. 6-7 weeks old female NOD/SCID mice were purchased from HFK Bio-Technology Co. Ltd. (Beijing, China). Each mouse was inoculated subcutaneously at the right flank region with 1×10^7^ HL-60 cells in 100 µl ice-cold PBS. The tumor-bearing mice were randomized into different groups with 6 mice per group. Treatment started when the mean tumor volume reached 56 mm^3^. 1.3 mg/kg of DN052 was sc administered once a week (qw), twice a week (biw) or three times a week (tiw), respectively. 1.3 mg/kg of motolimod was sc administered once a week. Vehicle treated mice were included as controls.

### Cytokine induction assays

To evaluate the activity of DN052 and motolimod in immune modulation, cytokine induction experiment was conducted in human peripheral blood mononuclear cells (PBMCs). Briefly, PBMCs were isolated from fresh human blood (Chempartner, Shanghai, China). 2×10^5^/100μL/well hPBMCs were plated on 96-well cell culture plates. Serial dilutions of compounds with concentration range from 2 nM to 30 µM were added in duplicate wells. LPS was included as a control. The plate was incubated in a 37°C, 5% CO_2_ incubator for 24 h. The supernatant was harvested for cytokine detection using MILLIPLEX MAP Human Cytokine/Chemokine Magnetic Bead Panel - Immunology Multiplex Assay (Millipore) following the manufacturer’s protocol. Cytokines including TNFα, IFNα2, IL-1α, IL-1β, IL-6, IL-8, IL-10, IL-12p40, IL-12p70, MIP-1α, MIP-1β, G-CSF and IFNγ were measured.

The cytokine induction activity of DN052 was further investigated in vivo using cynomolgus monkeys. Cynomolgus monkeys were purchased from Guangxi Xiongsen Primate Laboratory Animals Co., Ltd. (Guangxi, China) and three 3-4 years old male monkeys were used in each group. A single dose of DN052 was sc administered to monkeys at 1, 3 and 10 mg/kg, respectively. A single dose of motolimod was sc administered at 1 mg/kg. Blood samples were collected pre-dose, 6 and 24 h post-dose. Levels of cytokines including IFNγ, IL-10, IL-1β, IL-6, IL-8 MIP-1α, MIP-1β, G-CSF, IL-12p40, IL-12p70 and TNFα were measured using MSD ECL cytokine assay kit following the manufacturer’s protocol.

### Toxicology and safety pharmacology studies

Toxicity of DN052 was evaluated in rats and cynomolgus monkeys. First, Non-Good Laboratory Practice (Non-GLP) dose tolerability study was performed to find appropriate dose ranges. DN052 was sc administered to rats and monkeys and standard toxicology endpoints were evaluated. Then, GLP toxicology study was conducted. All aspects of the GLP studies were conducted in accordance with China Food and Drug Administration Good Laboratory Practice for Nonclinical Laboratory Studies and the United States Food and Drug Administration (FDA) Good Laboratory Practice (GLP) for Nonclinical Laboratory Studies. In the rat GLP study, DN052 was sc administered once a week for 3 weeks to Sprague Dawley rats at 1, 6 and 12 mg/kg, respectively, followed by a 4-week recovery phase. Vehicle treated rats were included as controls. 7 to 9 weeks old rats were purchased from Vital River Laboratory Animal Technology Co. Ltd. (Beijing, China) and 30 rats (15 males and 15 females) were used in each group. Assessment of toxicity was based on mortality, clinical observations, skin injection site reaction, body weight, food consumption, ophthalmic observations and clinical and anatomic pathology. In the monkey GLP study, DN052 was sc administered once a week for 3 weeks to cynomolgus monkeys at 0.3, 0.6 and 2 mg/kg, respectively, followed by a 4-week recovery phase. Vehicle treated monkeys were included as controls. 3 to 4 years old cynomolgus monkeys were purchased from Suzhou Xishan ZhongKe Laboratory Animal Co., Ltd. (Suzhou, Jiangsu, China) and 10 monkeys (5 males and 5 females) were used in each group. Assessment of toxicity was based on mortality, clinical observations, skin injection site reaction, body weight, food consumption, ophthalmology, vital signs (including body temperature, blood pressure and respiration rate), electrocardiography (ECG) and clinical and anatomic pathology. In the GLP safety pharmacology study, animals were monitored for cardiovascular function in cynomolgus monkeys administered with 0.3, 0.6 and 2 mg/kg of DN052.

### Statistical analyses

One-way ANOVA followed by Dunnett’s post-test and paired t-test were applied to assess the statistical significance of differences between treatment groups using GraphPad Prism. Statistical significance was accepted when *p* < 0.05.

## RESULTS

### DN052 was a novel, potent and selective TLR8 agonist

The *in vitro* activity of DN052 and motolimod was determined in cell-based assays (Fig. 1A). The EC_50_ value of DN052 in hTLR8 agonist activity was 6.7 nM. The EC_50_ value of the reference compound motolimod was 120.4 nM. The results indicated DN052 was about 16-fold more potent than motolimod in activating TLR8 *in vitro*. The CC_50_ values of DN052 and motolimod were 11,074 nM and 6,959 nM, respectively, suggesting both compounds did not have substantial non-specific cell killing activity (Fig. 1A). To evaluate the selectivity of DN052 for TLR8 over several other related TLRs including TLR4, TLR7 and TLR9, cell-based assays in HEK-Blue™ hTLR4, hTLR7 or hTLR9 cell line expressing the corresponding human TLR and SEAP reporter gene was used. The EC_50_ values of DN052 in the hTLR4, hTLR7 and hTLR9 agonist activity assays were greater than 50 μM, the highest concentration used in the assays, indicating DN052 was highly selective for TLR8 with no activity over TLR4, TLR7 or TLR9 (Fig. 1A). The EC_50_ values of motolimod in the hTLR4 and hTLR9 agonist activity assays were greater than 50 μM. The EC_50_ value of motolimod in the hTLR7 agonist activity assay was 12.1 μM indicating motolimod had weak activity over TLR7 in addition to its predominant agonistic activity for TLR8 (Fig. 1A).

**FIGURE 1.**
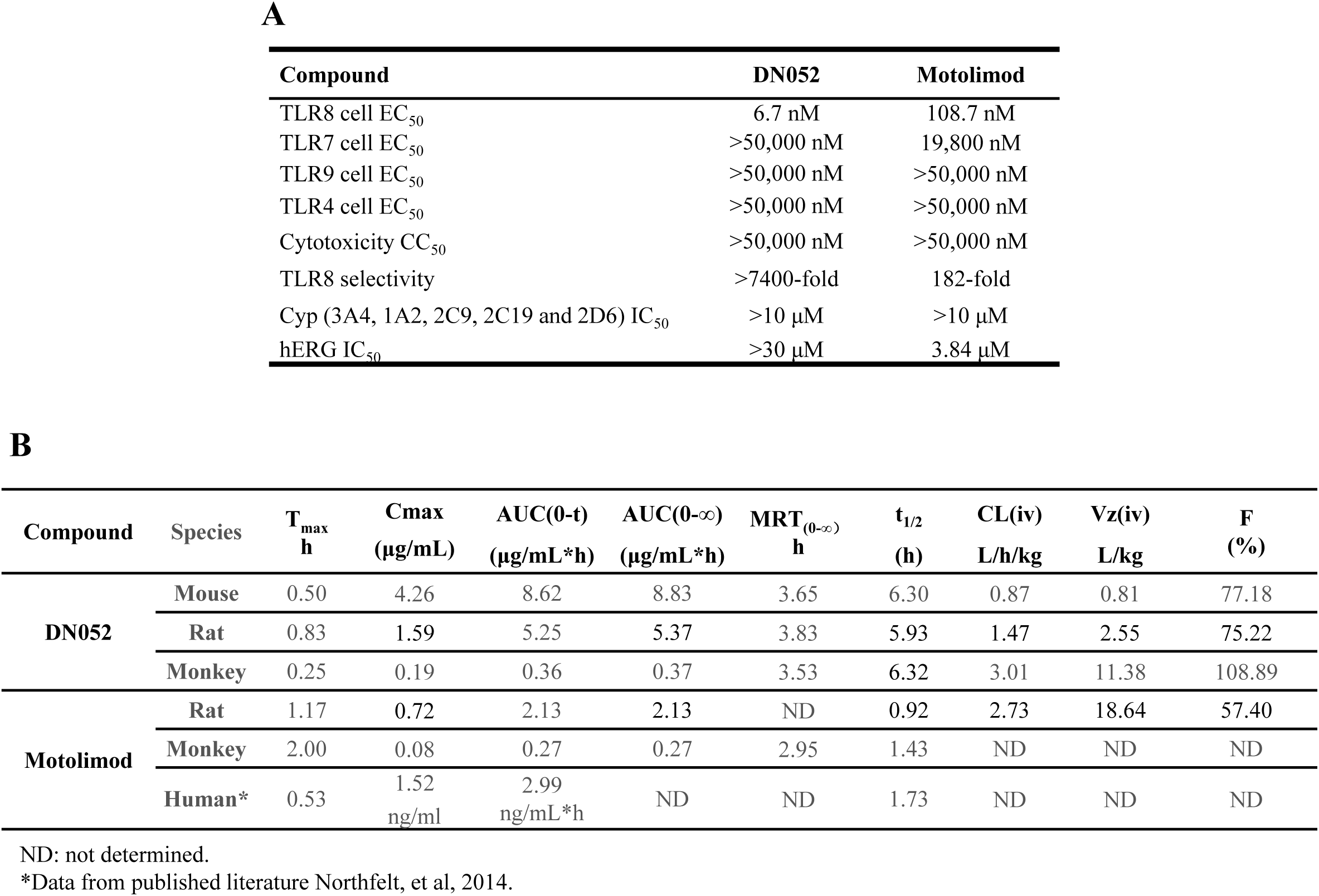
In vitro and DMPK profiles of DN052. (A) In vitro profiles of DN052. The chemical structure and synthesis of DN052 were described in the published patent WO/2017/190669. (B) Mouse PK: iv dose at 2.5 mg/kg and sc dose at 10 mg/kg for DN052. Rat PK: iv dose at 2.5 mg/kg and sc dose at 10 mg/kg for both motolimod and DN052. Monkey PK: iv dose at 1 mg/kg for DN052; sc dose at 1 mg/kg for both motolimod and DN052. Human PK: sc dose at 0.1 mg/m2 for motolimod. ND: not determined. *Data from published literature Northfelt, et al, 2014.

To assess potential off-target effects of DN052 in comparison to motolimod, the Cerep screen was conducted for DN052 and motolimod in receptor binding, enzyme and uptake assays involving 44 targets. Results showing an inhibition or stimulation higher than 50% are considered to represent significant effects of the test compounds. As shown in Supplemental Fig. 1, overall, DN052 and motolimod shared similar profiles with DN052 displaying fewer off-targets than motolimod. Motolimod showed 59.2% against potassium channel hERG suggesting motolimod might have potential cardiac toxicity at high concentration. In contrast, DN052 did not show any effect on the potassium hERG in the Cerep screen suggesting DN052 may be safer than motolimod when used at high concentration levels. Of note, this Cerep screen was performed with 10 μM compounds which was a very high concentration unlikely to be used in pharmacology studies. In addition, motolimod has advanced to clinical phase 2 trials without report of unacceptable adverse events (36, 37). Taken together, the off-target results supported further development of DN052.

### DN052 had favorable drug metabolism and pharmacokinetic profiles

DN052 showed clean CYP profile with IC50 over 10 μM for all major CYP isoenzymes tested including 3A4, 1A2, 2C9, 2C19 and 2D6 (Fig. 1A). DN052 had favorable hERG parameter with IC50 over 30 μM whereas motolimod’s hERG IC50 was 3.84 μM (Fig. 1A) suggesting motolimod may have potential cardiac toxicity liability when used at high doses. Interestingly, the Cerep off-target screen also showed motolimod had effect on hERG (Supplemental Fig. 1). The observation of motolimod’s hERG effect in two independent assays strongly suggested motolimod’s cardiac liability may limit its dose levels. Comparison of PK parameters with different dosing routes including iv, ip, sc and po in rats revealed that sc dosing was more desirable (Table I and Fig. 2) and therefore sc dosing was selected in subsequent studies. Furthermore, DN052 showed superior in vivo PK profile than motolimod in rats and monkeys (Fig. 1B). With sc administration, DN052 had higher AUC, Cmax, bioavailability and longer t_1/2_ than motolimod. The mouse PK of DN052 also showed favorable profiles (Fig. 1B).

**Table I.** Comparison of PK of DN052 with different dosing routes in rats.

| Animal ID | $t_{1/2}$ | $T_{max}$ | $C_{max}$ | $AUC_{(0-t)}$ | $AUC_{(0-\infty)}$ | $V_z$ | $CL_z$ | $MRT_{(0-\infty)}$ |
| --- | --- | --- | --- | --- | --- | --- | --- | --- |
|  | hr | hr | ng/L | ng/L*hr | ng/L*hr | L/kg | L/hr/kg | hr |
| **<u>IV (2.5 mg/kg)</u>** |  |  |  |  |  |  |  |  |
| 101 | 1.26 | 0.083 | 3790.58 | 1444.12 | 1446.82 | 3.15 | 1.73 | 0.46 |
| 102 | 1.28 | 0.083 | 6626.67 | 2156.96 | 2161.94 | 2.14 | 1.16 | 0.27 |
| 103 | 1.07 | 0.083 | 4242.69 | 1637.17 | 1642.71 | 2.36 | 1.52 | 0.42 |
| Mean | 1.21 | 0.083 | 4886.65 | 1746.08 | 1750.49 | 2.55 | 1.47 | 0.38 |
| SD | 0.12 | 0.000 | 1523.77 | 368.69 | 369.54 | 0.53 | 0.29 | 0.10 |
| Animal ID | $t_{1/2}$ | $T_{max}$ | $C_{max}$ | $AUC_{(0-t)}$ | $AUC_{(0-\infty)}$ | $MRT_{(0-\infty)}$ | F | |
|  | hr | hr | ng/L | ng/L*hr | ng/L*hr | hr | % |  |
| **<u>SC (10 mg/kg)</u>** |  |  |  |  |  |  |  |  |
| 201 | 5.13 | 0.500 | 1832.27 | 5400.93 | 5472.83 | 3.31 | 77.33 |  |
| 202 | 5.86 | 1.000 | 1544.70 | 4707.30 | 4823.60 | 3.95 | 67.40 |  |
| 203 | 6.79 | 1.000 | 1383.32 | 5653.51 | 5799.55 | 4.23 | 80.95 |  |
| Mean | 5.93 | 0.833 | 1586.76 | 5253.92 | 5365.33 | 3.83 | 75.22 |  |
| SD | 0.83 | 0.289 | 227.41 | 489.94 | 496.78 | 0.48 | 7.01 |  |
| Animal ID | $t_{1/2}$ | $T_{max}$ | $C_{max}$ | $AUC_{(0-t)}$ | $AUC_{(0-\infty)}$ | $MRT_{(0-\infty)}$ | F | |
|  | hr | hr | ng/L | ng/L*hr | ng/L*hr | hr | % |  |
| **<u>PO (10 mg/kg)</u>** |  |  |  |  |  |  |  |  |
| 301 | 0.56 | 2.000 | 928.73 | 2104.83 | 2105.91 | 2.27 | 30.14 |  |
| 302 | 0.48 | 4.000 | 374.47 | 916.24 | 916.99 | 3.73 | 13.12 |  |
| 303 | NA | 4.000 | 309.82 | 917.56 | NA | NA | 13.14 |  |
| Mean | 0.52 | 3.333 | 537.67 | 1312.88 | 1511.45 | 3.00 | 18.80 |  |
| SD | 0.06 | 1.155 | 340.20 | 685.85 | 840.69 | 1.03 | 9.82 |  |

**FIGURE 2.**
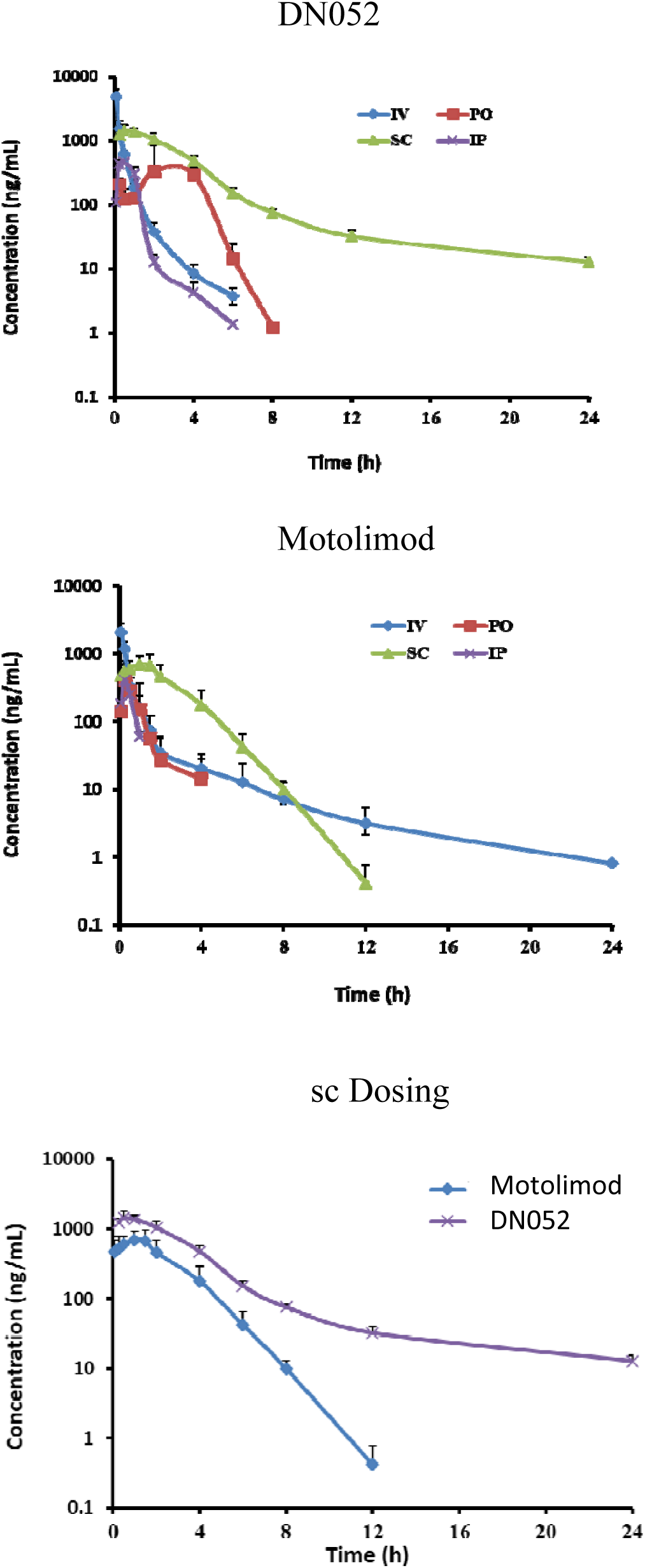
Comparison of PK with different dosing routes. DN052 and motolimod were iv, ip, sc or po administered to Sprague Dawley rats, respectively. Overall, sc appeared to have better PK than the other dosing routes for DN052. DN052 showed superior PK profile than motolimod.

### DN052 strongly inhibited tumor growth as a single agent or in combination with other anti-cancer drugs

TLR8 is unique among TLRs in that TLR8 manifests species specific responses to its ligands with much diminished activity in rodents (42), thereby limiting the usefulness of commonly used mouse models in evaluating the in vivo biological activities of TLR8 agonists. This limitation has been a major hurdle in drug discovery effort targeting TLR8. We addressed this limitation by using two different approaches: 1) Immune-competent mouse syngeneic tumor models: high doses of DN052 were used to offset the low TLR8 activity in rodents; 2) Human AML mouse xenograft model: human HL-60 AML cells could be directly targeted by TLR8 agonist through its effect of inducing terminal differentiation-mediated tumor suppression (25).

DN052 strongly suppressed tumor growth in a dose dependent manner as a single agent when used at high doses at 40, 80 and 160 mg/kg in immune-competent CT-26 mouse syngeneic colon cancer model (Fig. 3A). DN052 was well-tolerated in the mice at all the doses tested (Fig. 3B). Similar result was observed in the immune-competent EMT6 mouse syngeneic breast cancer model in which DN052 markedly suppressed tumor growth as a single agent and resulted in complete tumor regression in 1/8, 2/8 and 3/8 tumor-bearing mice at 40, 80 and 160 mg/kg, respectively (Fig. 3C, 3D). Moreover, when evaluated side-by-side under the same condition, DN052 appeared to be more efficacious than motolimod in EMT6 model (Fig. 3C). The strong efficacy produced by DN052 single agent suggested that DN052 had potential to be used as monotherapy in treating cancer patients.

**FIGURE 3.**
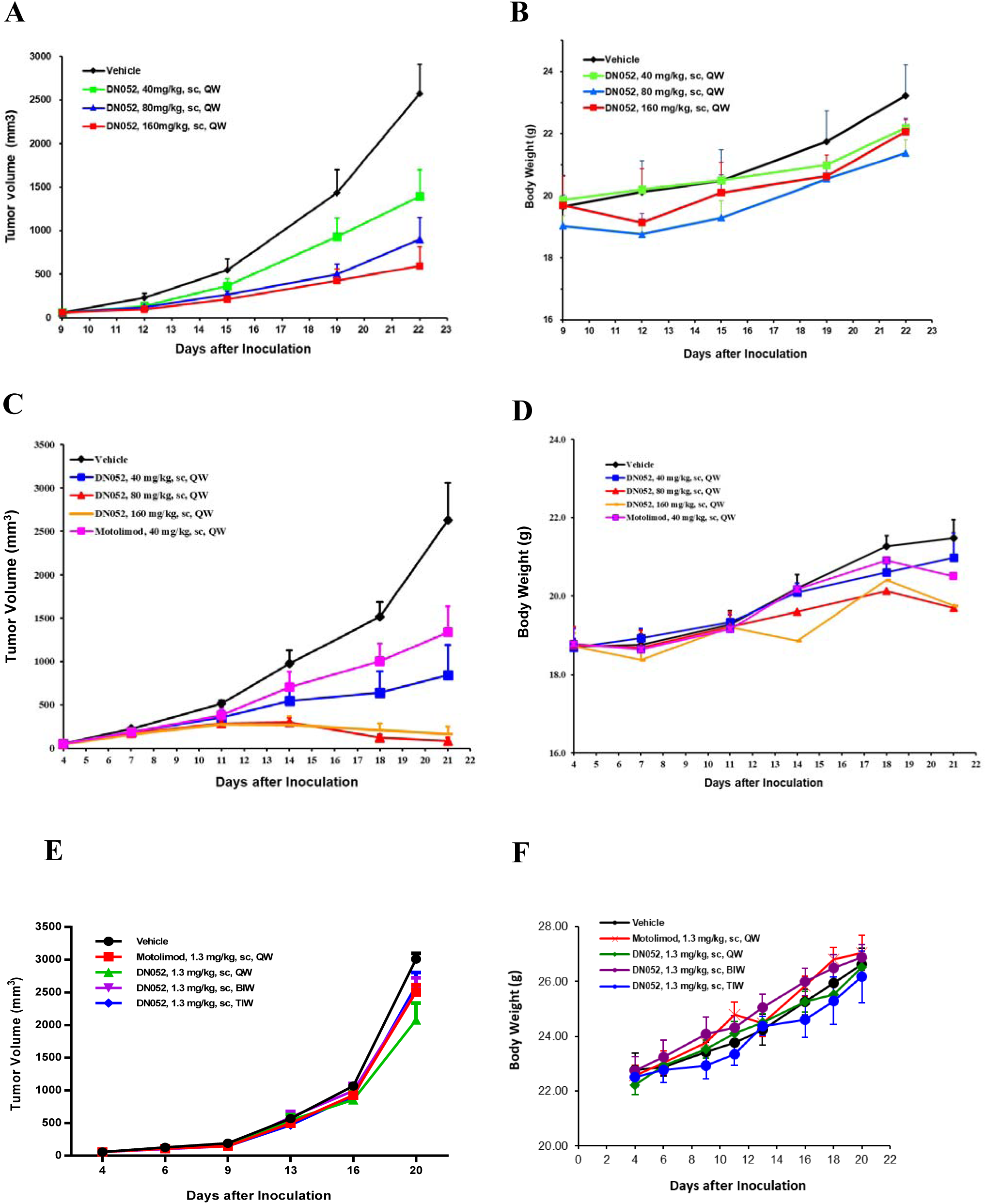
DN052 was more efficacious than motolimod and well-tolerated as a single agent in various mouse tumor models. DN052 treatment resulted in significant tumor growth inhibition in colon cancer (A, B), breast cancer (C, D) and AML (E, F) models. DN052 more strongly suppressed tumor growth than motolimod under the same conditions.

Furthermore, to explore its application in combination therapies addressing critical unmet medical needs, we carried out a series of combination studies in several cancer models. Combination of DN052 and the chemotherapeutic drug cyclophosphamide (CTX) resulted in stronger tumor suppression than either agent alone in immune-competent CT26 mouse syngeneic colon cancer model (Fig. 4A). In another combination study, while EMT6 model was largely resistant to AZD-1775, a WEE1 inhibitor currently in phase 2 clinical development (43), addition of DN052 drastically enhanced tumor growth inhibition compared to either agent alone (Fig. 4B). Similarly, combination of DN052 and AZD-1775 increased efficacy in immune-competent H22 mouse syngeneic liver cancer model (Fig. 4C). Moreover, addition of DN052 increased tumor growth inhibition produced by sorafenib, a standard of care (SOC) agent for liver cancer in H22 model (Fig. 4C). DN052 or anti-PD-1 monoclonal antibody (αPD-1) suppressed tumor growth as single agent in immune-competent MC-38 mouse syngeneic colon cancer model (Fig. 4D). Combination of DN052 and αPD-1 exhibited stronger tumor suppression than either agent alone though the effect of combination appeared less pronounced based on tumor growth inhibition rates (Fig. 4D). However, it is noteworthy that combination of DN052 and αPD-1 resulted in complete tumor regression in 1/7 MC-38 tumor-bearing mice whereas no complete tumor regression in mice treated with single agents further suggesting DN052 enhanced αPD-1’s anti-cancer effect (Fig. 4D).

**FIGURE 4.**
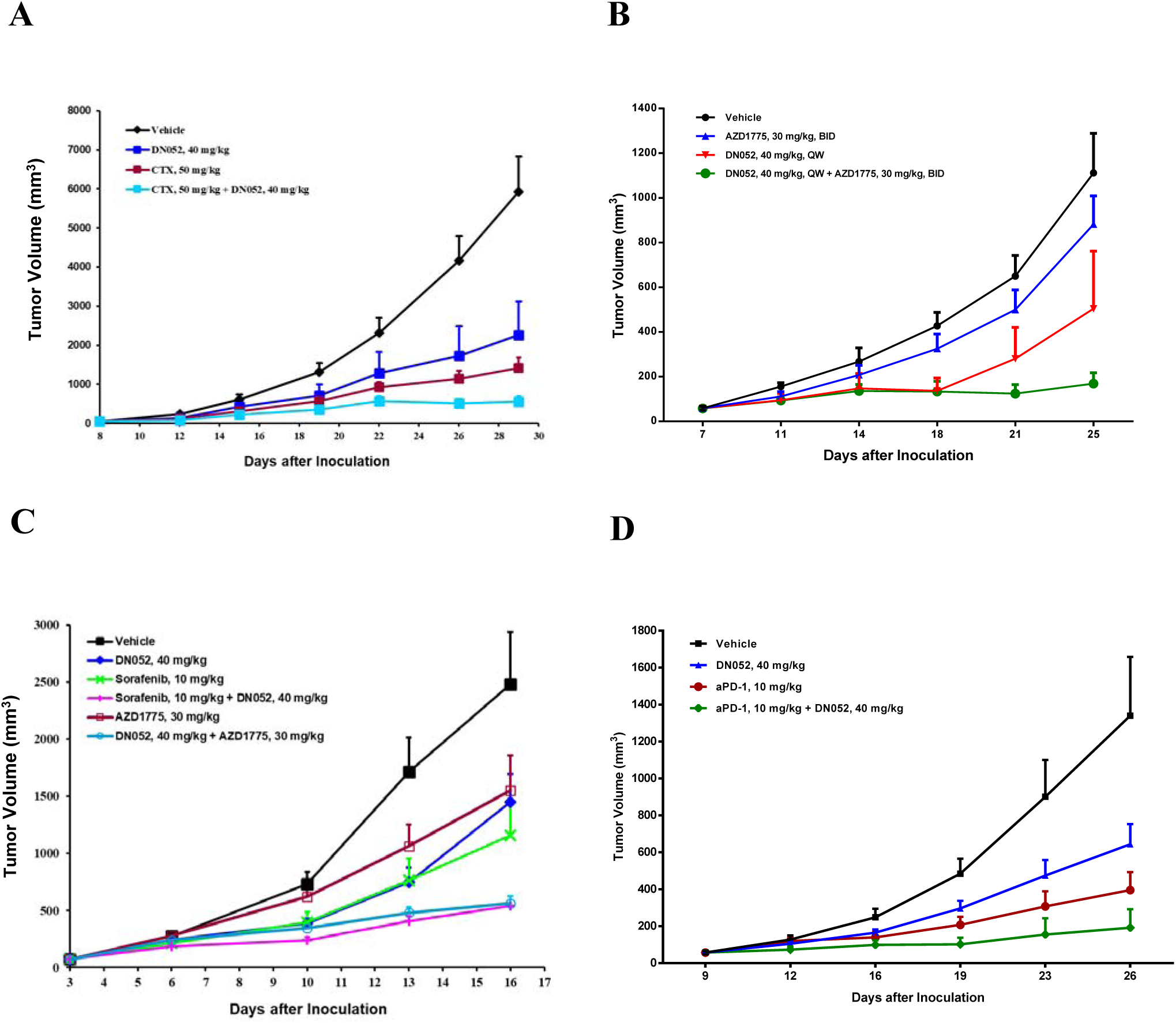
DN052 further reduced tumor growth when combined with other anti-cancer agents in mouse tumor models. Combination of DN052 and cyclophosphamide, WEE1 inhibitor AZD-1775, sorafenib or αPD-1 increased tumor suppression in several mouse tumor models: CT26 model (A), EMT6 model (B), H22 model (C) and MC38 model (D).

Although the in vivo efficacy observed in the mouse syngeneic models was remarkable, the high doses used in these studies are less predictive of the human dose in clinical trials because of TLR8’s species specificity. To gain further information about the in vivo efficacious doses more relevant to humans, we conducted a study using human HL-60 AML mouse xenograft model. DN052 impeded tumor growth in the immune-deficient mice bearing human HL-60 AML when dosed sc at 1.3 mg/kg. DN052 caused stronger tumor growth inhibition than motolimod under the same condition with tumor growth inhibition (TGI) rate 31% vs 17% indicating DN052 was more active in vivo than motolimod (Fig. 3E). The stronger activity of DN052 than motolimod was consistent with the results in the immune competent EMT6 mouse syngeneic model as well as the in vitro cell based assays. Furthermore, 1.3 mg/kg QW produced stronger efficacy than BIW and TIW indicating that infrequent dosing can be achieved. Importantly, DN052 appeared to be efficacious with all the dosing schedules tested. During the study, there was no mortality or significant changes in mouse body weight between test compounds-treated animals and the vehicle controls indicating all the treatment was well-tolerated by the animals (Fig. 3F). Based on these results and the published literature reporting QW dosing for motolimod (37), QW dosing schedule was chosen for DN052 in both efficacy and toxicology studies.

Of note, since the HL-60 xenograft model used in this study was immune-deficient, the anti-tumor activity of DN052 and motolimod observed was unlikely mediated through immune response. The direct effect on the human HL-60 cells through induction of leukemia cell terminal differentiation accounted for the anti-tumor effect which is supported by the literature (25). The relatively modest tumor growth inhibition observed in this model could be underestimated given the immune-deficient background where the TLR8’s immune modulation was largely bypassed in this model.

### DN052 induced pro-inflammatory cytokines

To understand the underlying mechanism for DN052’s anticancer effect, an *ex vivo* human PBMC assay was used to evaluate the immune modulatory activity of DN052. Motolimod was included as a reference compound. As shown in Table II, DN052 strongly induced the pro-inflammatory cytokines including TNFα, IFNα_2_, IL-1α, IL-1β, IL-6, IL-8, IL-10, IL-12p40, IL-12p70, MIP-1α, MIP-1β, G-CSF and IFN γ. Overall, DN052 was more potent in inducing the cytokines than motolimod. Several of the cytokines showed robust induction for both DN052 and motolimod including MIP-1β, MIP-1α, IL-1β, IL-6, IL-8, IL-12p40 and TNFα. Comparison of the EC_50_ values for these cytokines between DN052 and motolimod indicated that DN052 was about 7-22 times more potent than motolimod.

**Table II.**
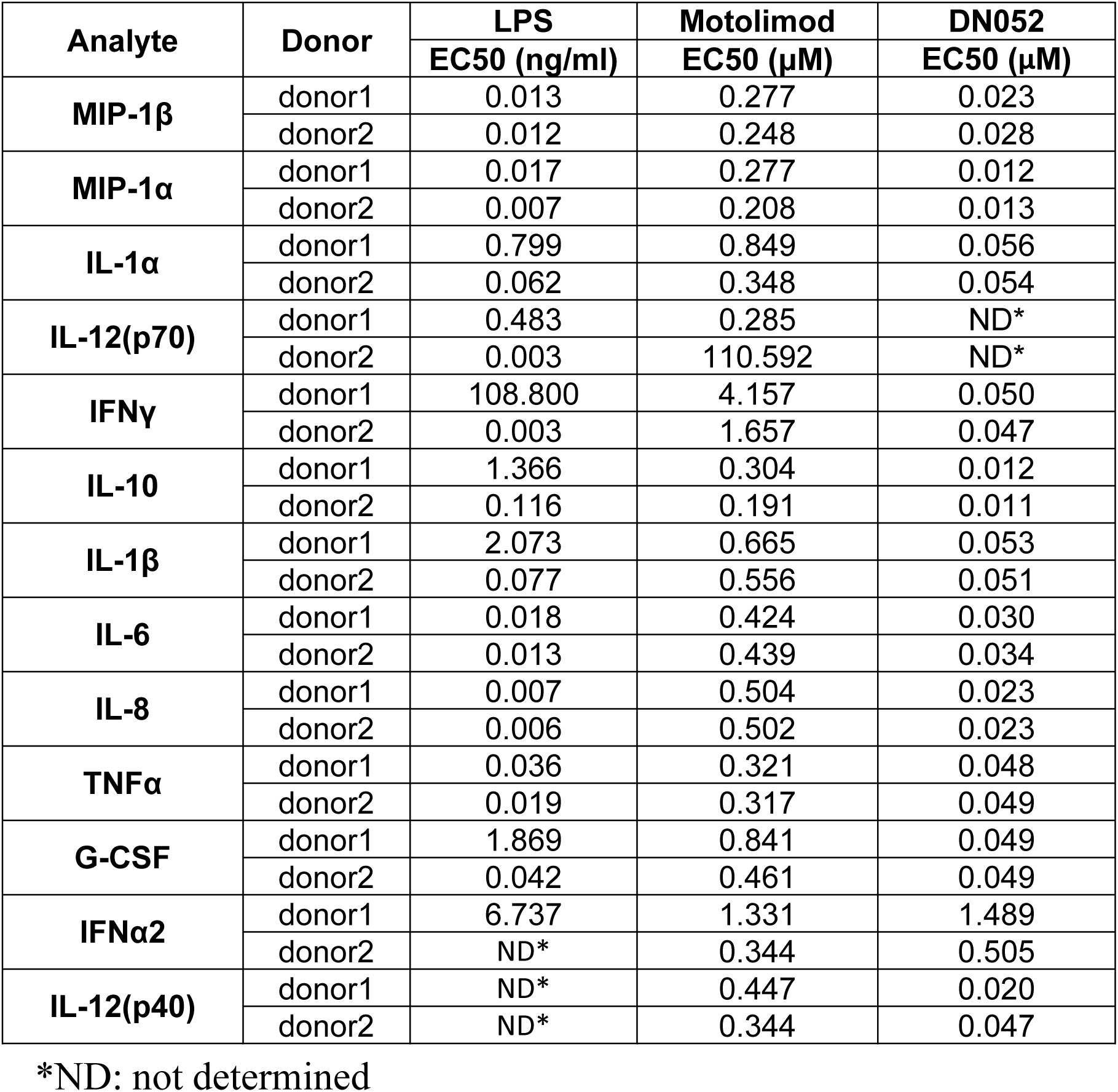
DN052 induced cytokines in ex vivo human PMBCs.

To further investigate the immune response to DN052 *in vivo*, DN052 was sc administered to cynomolgus monkeys and changes in serum levels of cytokines IFNγ, IL-1β, IL-6, IL-8, IL-10, IL-12P40, IL-12P70, TNFα, G-CSF, MIP-1α, MIP-1β were measured. Consistent with the hPBMC result, treatment with DN052 resulted in strong induction of the cytokines. DN052 was more effective than motolimod when used at the same dose 1 mg/kg (Table III).

**Table III.** DN052 induced cytokines in monkeys.

| PD Marker | Timepoint (h) | Mean Values (pg/mL) |  |  |  |
| --- | --- | --- | --- | --- | --- |
|  |  | Motolimod (1 mg/kg, sc) | DN052 (1 mg/kg, sc) | DN052 (3 mg/kg, sc) | DN052 (10 mg/kg, sc) |
| IFN-γ | 0 | 1.017 | 0.785 | 1.249 | 0.397 |
|  | 6 | 0.500 | 39.126 | 9.894 | 9.544 |
|  | 24 | 0.462 | 1.062 | 0.000 | 1.209 |
| IL-10 | 0 | 0.271 | 0.295 | 0.366 | 0.217 |
|  | 6 | 15.222 | 726.647 | 680.726 | 1497.839 |
|  | 24 | 1.347 | 5.166 | 6.821 | 52.473 |
| IL-1β | 0 | 0.000 | 0.130 | 0.431 | 0.000 |
|  | 6 | 0.286 | 2.196 | 3.085 | 4.071 |
|  | 24 | 0.099 | 0.231 | 0.853 | 0.456 |
| IL-6 | 0 | 1.510 | 3.403 | 7.566 | 2.574 |
|  | 6 | 342.661 | 2747.061 | 1850.460 | 9266.511 |
|  | 24 | 12.185 | 120.795 | 195.354 | 1464.089 |
| IL-8 | 0 | 0.000 | 0.257 | 1.085 | 0.238 |
|  | 6 | 0.378 | 1.215 | 1.964 | 5.416 |
|  | 24 | 0.119 | 0.401 | 0.716 | 1.044 |
| MIP-1α | 0 | 71.636 | 56.670 | 58.578 | 65.268 |
|  | 6 | 227.638 | 492.577 | 487.888 | 594.484 |
|  | 24 | 87.417 | 152.799 | 151.320 | 181.215 |
| MIP-1β | 0 | 93.266 | 215.299 | 195.431 | 210.978 |
|  | 6 | 1655.094 | 3190.677 | 1934.886 | 6215.896 |
|  | 24 | 221.881 | 659.556 | 692.643 | 1158.540 |
| IL-12P40 | 0 | 131.413 | 191.908 | 279.654 | 158.623 |
|  | 6 | 305.212 | 7295.739 | 5927.984 | 5525.537 |
|  | 24 | 118.409 | 653.428 | 553.852 | 633.985 |
| G-CSF | 0 | 7.135 | 29.193 | 41.588 | 29.765 |
|  | 6 | 345.154 | 2411.265 | 2081.330 | 6683.544 |
|  | 24 | 32.168 | 58.352 | 127.944 | 576.160 |
| IL-12P70 | 0 | 2.582 | 2.458 | 2.214 | ND* |
|  | 6 | 5.154 | 9.084 | 2.977 | ND* |
|  | 24 | 3.305 | 2.834 | 0.000 | ND* |
| TNF-α | 0 | ND* | ND* | 1.931 | 3.033 |
|  | 6 | ND* | ND* | 2.684 | 3.380 |
|  | 24 | ND* | ND* | 1.676 | 2.501 |

\*ND: not determined

### DN052 had favorable safety profiles

The potential toxicity of DN052 when administered once weekly via subcutaneous injection to rodents (rats) or non-rodent large animals (monkeys) were investigated. In the Sprague Dawley rat GLP study, DN052 was dosed at 2, 6, 12 mg/kg, sc, QW, respectively. 12 mg/kg was determined to be no observed adverse effect level (NOAEL). Mild skin injection site reaction was observed. Maximum tolerated dose (MTD) was not reached. The strong tolerability exhibited by rats treated with DN052 was expected given TLR8’s species specificity in that TLR8’s activity in rodents is very low (42). In contrast, in the cynomolgus monkey GLP study, DN052 was dosed at 0.3, 0.6 and 2 mg/kg, sc, QW, respectively. Skin injection site reaction was observed at all dose levels. Microscopic examination at the terminal sacrifice showed minimal to slight acute or chronic-active inflammation, mixed cell infiltrates, and/or aggregation of foamy macrophages in the subcutis of animals administered 0.3 and 0.6 mg/kg and ulcer and/or acute inflammation in the lumbar skin/subcutis of animals administered 2.0 mg/kg. Microscopic changes at the injection sites of animals administered 0.3 or 0.6 mg/kg were considered non-adverse. At the recovery sacrifice, these microscopic findings had completely recovered indicating that the skin injection site reaction was reversible. 0.6 mg/kg was determined to be NOAEL and the highest non-severely toxic dose (HNSTD) is between 0.6 mg/kg and 2.0 mg/kg. No DN052-related changes to blood pressure were noted. No abnormal ECG waveforms or arrhythmias attributed to DN052 were observed in the cynomolgus monkeys administered 0.3, 0.6, or 2 mg/kg.

As reported in the literature, the highest dose of motolimod used in phase 1 clinical trials was 3.9 mg/m^2^ (40) and its phase 2 doses were 2.5-3.5 mg/m^2^ (34). 3.9 mg/m^2^ in humans is equivalent to approximately 0.3 mg/kg in monkeys, which was half of the NOAEL of DN052 (0.6 mg/kg). Therefore, the toxicity result indicated that DN052 had favorable safety profile. On the other hand, the in vitro cell based assays showed that DN052 was 16-fold more potent than motolimod. The in vivo efficacy study in tumor models demonstrated that DN052 was more active than motolimod. Taken together with the toxicity data, these results suggested that DN052 may have larger therapeutic index than motolimod in humans.

## DISCUSSION

Harnessing the host’s immune system to eradicate cancer cells has become a powerful new approach to cancer therapy in recent years partly attributing to the unprecedented clinical success of immune checkpoint inhibitors (2, 3). However, despite the significant progress in cancer immunotherapy with checkpoint blockade, majority of cancer patients do not respond to the current therapy (4). The lack of response to immune checkpoint blockade in majority of cancer patients represents an urgent unmet medical need. Human body’s immune system consists of innate and adaptive immune pathways (1). While releasing the immune checkpoint blockage in the adaptive immunity has proved effective therapy, targets involved in the innate immunity are just beginning to show promise in the fight against cancer (4). We hypothesized that targeting innate immunity would enhance the anticancer efficacy produced by drugs targeting the adaptive immunity because of their complementary nature as host’s immune defense system.

TLR8 was chosen because it is one of the most important molecules of the innate immunity (42, 44-46). Accumulating evidence indicated that activation of TLR8 could reverse Treg and MDSC mediated immune suppression resulting in strong tumor inhibition (16, 19-21). One of the major causes of cancer immunotherapy failure is potent suppression of immune response by Treg or MDSC cells (21-23). Therefore, TLR8 agonists possess the potential to turn immune unresponsive “cold” tumors to immune responsive “hot” tumors, thereby addressing the urgent unmet medical need in tumor immunotherapy.

DN052 is a novel TLR8 selective agonist displaying differentiated profiles compared to motolimod in that DN052 specifically targets TLR8 while sparing TLR7. Several TLR agonists including TLR4, 7, TLR7/8 dual and TLR9 agonists can be used mainly as a topical drug or dosed locally such as intratumoral injection because they are too toxic if used systemically (5). This has been a major hurdle which limits their application in the clinic (13, 32, 33). In contrast, our study demonstrated that the high selectivity of DN052 on TLR8 showed better safety allowing systemic dosing which was well tolerated in the in vivo studies. To the best of our knowledge, DN052 is the first truly selective TLR8 agonist. DN052 also showed stronger activity than motolimod in the cell based assays and superior DMPK profiles in rats and monkeys and excellent PK in mice. TLR8 was previously thought to be non-functional in mice (42). However, more recent studies indicated that TLR8 plays crucial roles in mice albeit its receptor activity is much diminished in rodents including mice and rats compared to other species such as humans (47). The species specificity of TLR8 hindered much of the in vivo studies in the context of drug discovery. To address this challenge, we applied high doses of DN052 in syngeneic mouse tumor models to offset the reduced receptor activity of TLR8 in mice. DN052 showed strong in vivo efficacy when used as a single agent in immune-competent mouse syngeneic tumor models for multiple cancer types.

To understand the mechanism for its tumor suppression, ex vivo human PBMC and in vivo monkey studies were carried out and DN052 activated immune response evidenced by strong induction of the proinflammatory cytokines including TNFα, IFNα_2_, IL-1α, IL-1β, IL-6, IL-8, IL-10, IL12p40, IL-12p70, MIP-1α, MIP-1β, G-CSF and INFγ. Overall, DN052 was more potent than motolimod in inducing the cytokines including INFγ, IL-12p40, TNFα which were reported to be predictive of clinical benefit in motolimod-treated cancer patients (36) suggesting DN052 may have superior efficacy in humans.

In another study, immune-deficient mouse xenograft model carrying human HL-60 AML was used to test if DN052 could inhibit tumor growth in vivo at lower doses. In this model, DN052 exerted the anticancer activity by directly targeting the human AML cells through induction of terminal differentiation-mediated tumor suppression (25). The result showed that treatment with low dose of DN052 resulted in significant tumor inhibition in the human AML mouse xenograft model. This approach overcame the species limitation seen in the syngeneic mouse tumor models where very high doses of DN052 had to be used because lower doses didn’t produce anticancer efficacy due to the much diminished TLR8 receptor activity in mice. Furthermore, the result suggested that unlike the high doses of DN052 used in syngeneic mouse models, low doses of DN052 are expected to be efficacious in humans.

Collectively, the series of in vivo efficacy studies using various mouse models representing different cancer types suggested that DN052 may be useful for both solid tumors and hematological malignancies in the clinic.

After demonstrating DN052 was efficacious when used as a single agent, we went on to ask if combination of DN052 with another anticancer agent would enhance the efficacy of single agents. Interestingly, several agents including cyclophosphamide, sorafenib, WEE1 inhibitor AZD1775 and αPD-1 monoclonal antibody showed increased tumor growth inhibition when combined with DN052. Although the precise mechanism underlying the enhanced efficacy is not fully understood and awaits further investigation, some of the mechanistic insight can be deduced from the literature. For example, the increased tumor growth inhibition produced by combining DN052 and the chemotherapeutic cyclophosphamide was likely due to the increased tumor mutation load caused by alkylating DNA. In addition, earlier reports suggested that cyclophosphamide treatment could result in mobilization of immune cells which might also play a role in increasing the anticancer efficacy (48). Combination of DN052 and WEE1 inhibitor showed strong synergy in suppressing tumor growth. The markedly enhanced efficacy was likely due to the enhanced antigen presentation as a result of higher tumor mutation load considering the function of WEE1 in DNA damage and repair (49). The combination therapy data of DN052 and αPD-1 supported the hypothesis that targeting both innate and adaptive immunity could increase anticancer efficacy. Taken together, the in vivo efficacy results strongly suggested that DN052 has the potential to be used as a backbone either as a single agent or in combination therapy in the clinic.

In clinical trials, motolimod is also administered subcutaneously and it has been reported that the skin injection site reaction (ISR) was correlated with significantly longer survival in terms of progression free survival (PFS) and overall survival (OS) in both ovarian, and head and neck cancer patients (36, 37). In preclinical studies, DN052 caused ISR more consistently than motolimod under the same conditions in animal studies suggesting DN052 may lead to better clinical outcome in human patients and ISR could be used as a surrogate marker in DN052 clinical trials. In addition, human papillomavirus (HPV)-positive status was shown to correlate with longer survival in head and neck cancer patients treated with motolimod (37). The high prevalence of HPV-positive head and neck cancer patients represents significant clinical indication. Therefore, HPV status may be useful in DN052 clinical trials.

DN052 is currently advancing in phase 1 trials in cancer patients in the US (ClinicalTrials.gov Identifier: NCT03934359) and both the safety and efficacy will be rigorously tested in the clinic. It is noteworthy that besides cancer indications, TLR8 agonists such as DN052 can be used for other indications including cancer vaccines (50) and the treatment of viral infections including HBV.

## DISCLOSURES

The authors have no financial conflicts of interest.

---

**Supplemental Figure 1.**
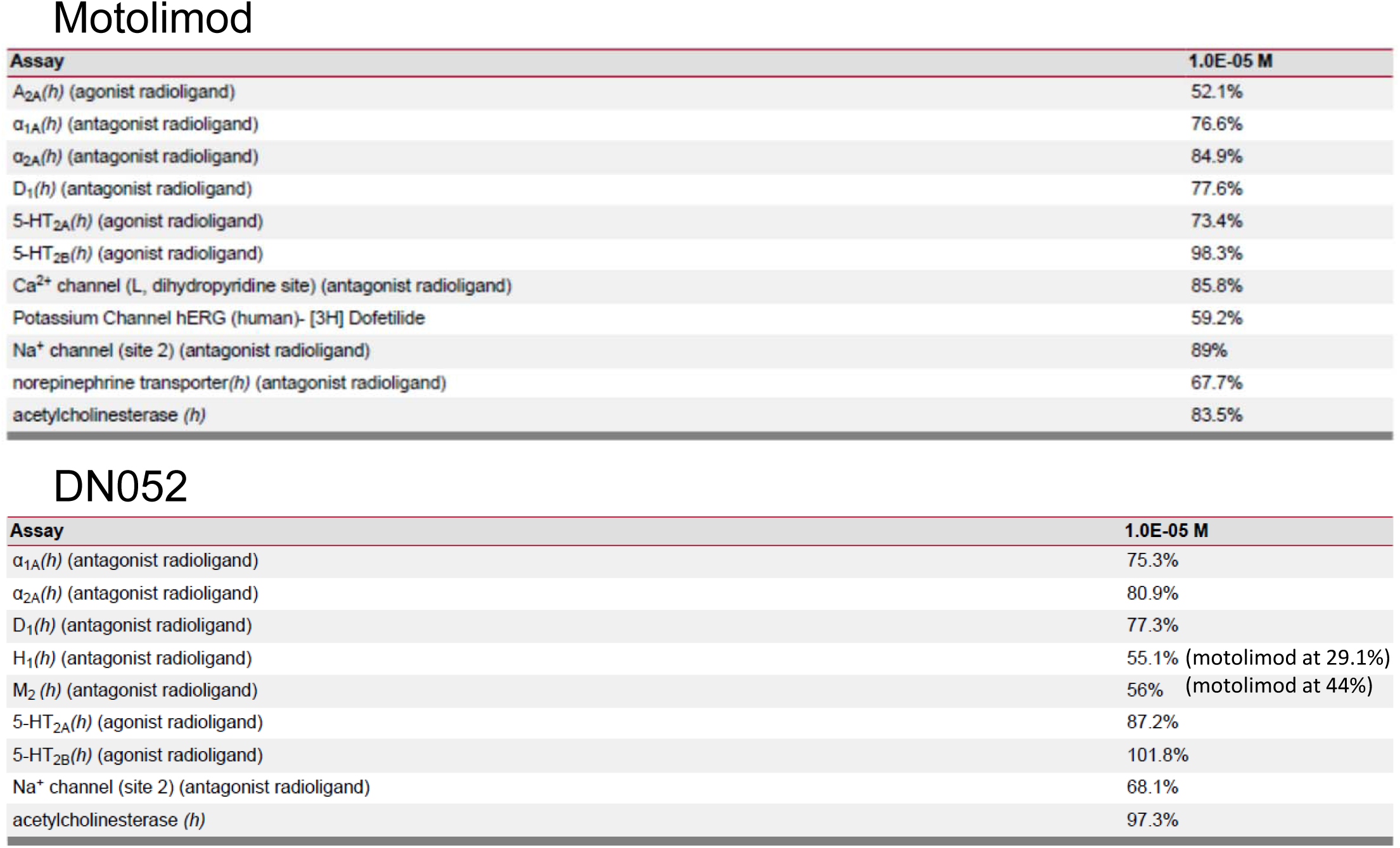
Cerep Screen Showed Less Off-target Effects of DN052 Compared to Motolimod.

## REFERENCES

1. Dranoff G. 2004. Cytokines in cancer pathogenesis and cancer therapy. Nat Rev Cancer 4: 11–22

2. Sharma P, Allison JP. 2015. The future of immune checkpoint therapy. Science 348: 56–61

3. Hahn AW, Gill DM, Pal SK, Agarwal N. 2017. The future of immune checkpoint cancer therapy after PD-1 and CTLA-4. Immunotherapy 9: 681–92

4. Demaria O, Cornen S, Daeron M, Morel Y, Medzhitov R, Vivier E. 2019. Harnessing innate immunity in cancer therapy. Nature 574: 45–56

5. Bourquin C, Pommier A, Hotz C. 2019. Harnessing the immune system to fight cancer with Toll-like receptor and RIG-I-like receptor agonists. Pharmacol Res: 104192

6. Shanker A, Marincola FM. 2011. Cooperativity of adaptive and innate immunity: implications for cancer therapy. Cancer Immunol Immunother 60: 1061–74

7. Pradere JP, Dapito DH, Schwabe RF. 2014. The Yin and Yang of Toll-like receptors in cancer. Oncogene 33: 3485–95

8. Seya T, Shime H, Ebihara T, Oshiumi H, Matsumoto M. 2010. Pattern recognition receptors of innate immunity and their application to tumor immunotherapy. Cancer Sci 101: 313–20

9. de Marcken M, Dhaliwal K, Danielsen AC, Gautron AS, Dominguez-Villar M. 2019. TLR7 and TLR8 activate distinct pathways in monocytes during RNA virus infection. Sci Signal 12

10. Bhatia S, Miller NJ, Lu H, Longino NV, Ibrani D, Shinohara MM, Byrd DR, Parvathaneni U, Kulikauskas R, Ter Meulen J, Hsu FJ, Koelle DM, Nghiem P. 2019. Intratumoral G100, a TLR4 Agonist, Induces Antitumor Immune Responses and Tumor Regression in Patients with Merkel Cell Carcinoma. Clin Cancer Res 25: 1185–95

11. Schon MP, Schon M. 2008. TLR7 and TLR8 as targets in cancer therapy. Oncogene 27: 190–9

12. Kuznik A, Panter G, Jerala R. 2010. Recognition of nucleic acids by Toll-like receptors and development of immunomodulatory drugs. Curr Med Chem 17: 1899–914

13. Huen AO, Rook AH. 2014. Toll receptor agonist therapy of skin cancer and cutaneous T-cell lymphoma. Curr Opin Oncol 26: 237–44

14. Krieg AM. 2007. Development of TLR9 agonists for cancer therapy. J Clin Invest 117: 1184–94

15. Komura F, Okuzumi K, Takahashi Y, Takakura Y, Nishikawa M. 2020. Development of RNA/DNA Hydrogel Targeting Toll-Like Receptor 7/8 for Sustained RNA Release and Potent Immune Activation. Molecules 25

16. Peng G, Guo Z, Kiniwa Y, Voo KS, Peng W, Fu T, Wang DY, Li Y, Wang HY, Wang RF. 2005. Toll-like receptor 8-mediated reversal of CD4+ regulatory T cell function. Science 309: 1380–4

17. Wang RF. 2006. Functional control of regulatory T cells and cancer immunotherapy. Semin Cancer Biol 16: 106–14

18. Wang RF, Miyahara Y, Wang HY. 2008. Toll-like receptors and immune regulation: implications for cancer therapy. Oncogene 27: 181–9

19. Li L, Liu X, Sanders KL, Edwards JL, Ye J, Si F, Gao A, Huang L, Hsueh EC, Ford DA, Hoft DF, Peng G. 2019. TLR8-Mediated Metabolic Control of Human Treg Function: A Mechanistic Target for Cancer Immunotherapy. Cell Metab 29: 103–23 e5

20. Liu X, Li L, Peng G. 2019. TLR8 reprograms human Treg metabolism and function. Aging (Albany NY) 11: 6614–5

21. Dang Y, Rutnam ZJ, Dietsch G, Lu H, Yang Y, Hershberg R, Disis ML. 2018. TLR8 ligation induces apoptosis of monocytic myeloid-derived suppressor cells. J Leukoc Biol 103: 157–64

22. Kiniwa Y, Miyahara Y, Wang HY, Peng W, Peng G, Wheeler TM, Thompson TC, Old LJ, Wang RF. 2007. CD8+ Foxp3+ regulatory T cells mediate immunosuppression in prostate cancer. Clin Cancer Res 13: 6947–58

23. Ye J, Ma C, Hsueh EC, Eickhoff CS, Zhang Y, Varvares MA, Hoft DF, Peng G. 2013. Tumor-derived gammadelta regulatory T cells suppress innate and adaptive immunity through the induction of immunosenescence. J Immunol 190: 2403–14

24. Wang RF. 2008. CD8+ regulatory T cells, their suppressive mechanisms, and regulation in cancer. Hum Immunol 69: 811–4

25. Ignatz-Hoover JJ, Wang H, Moreton SA, Chakrabarti A, Agarwal MK, Sun K, Gupta K, Wald DN. 2015. The role of TLR8 signaling in acute myeloid leukemia differentiation. Leukemia 29: 918–26

26. Slade HB, Owens ML, Tomai MA, Miller RL. 1998. Imiquimod 5% cream (Aldara). Expert Opin Investig Drugs 7: 437–49

27. Stary G, Bangert C, Tauber M, Strohal R, Kopp T, Stingl G. 2007. Tumoricidal activity of TLR7/8-activated inflammatory dendritic cells. J Exp Med 204: 1441–51

28. Adams S, Kozhaya L, Martiniuk F, Meng TC, Chiriboga L, Liebes L, Hochman T, Shuman N, Axelrod D, Speyer J, Novik Y, Tiersten A, Goldberg JD, Formenti SC, Bhardwaj N, Unutmaz D, Demaria S. 2012. Topical TLR7 agonist imiquimod can induce immune-mediated rejection of skin metastases in patients with breast cancer. Clin Cancer Res 18: 6748–57

29. Michaelis KA, Norgard MA, Zhu X, Levasseur PR, Sivagnanam S, Liudahl SM, Burfeind KG, Olson B, Pelz KR, Angeles Ramos DM, Maurer HC, Olive KP, Coussens LM, Morgan TK, Marks DL. 2019. The TLR7/8 agonist R848 remodels tumor and host responses to promote survival in pancreatic cancer. Nat Commun 10: 4682

30. Geller MA, Cooley S, Argenta PA, Downs LS, Carson LF, Judson PL, Ghebre R, Weigel B, Panoskaltsis-Mortari A, Curtsinger J, Miller JS. 2010. Toll-like receptor-7 agonist administered subcutaneously in a prolonged dosing schedule in heavily pretreated recurrent breast, ovarian, and cervix cancers. Cancer Immunol Immunother 59: 1877–84

31. Bergmann JF, de Bruijne J, Hotho DM, de Knegt RJ, Boonstra A, Weegink CJ, van Vliet AA, van de Wetering J, Fletcher SP, Bauman LA, Rahimy M, Appleman JR, Freddo JL, Janssen HL, Reesink HW. 2011. Randomised clinical trial: anti-viral activity of ANA773, an oral inducer of endogenous interferons acting via TLR7, in chronic HCV. Aliment Pharmacol Ther 34: 443–53

32. Fakhari A, Nugent S, Elvecrog J, Vasilakos J, Corcoran M, Tilahun A, Siebenaler K, Sun J, Subramony JA, Schwarz A. 2017. Thermosensitive Gel-Based Formulation for Intratumoral Delivery of Toll-Like Receptor 7/8 Dual Agonist, MEDI9197. J Pharm Sci 106: 2037–45

33. Mullins SR, Vasilakos JP, Deschler K, Grigsby I, Gillis P, John J, Elder MJ, Swales J, Timosenko E, Cooper Z, Dovedi SJ, Leishman AJ, Luheshi N, Elvecrog J, Tilahun A, Goodwin R, Herbst R, Tomai MA, Wilkinson RW. 2019. Intratumoral immunotherapy with TLR7/8 agonist MEDI9197 modulates the tumor microenvironment leading to enhanced activity when combined with other immunotherapies. J Immunother Cancer 7: 244

34. Chow LQM, Morishima C, Eaton KD, Baik CS, Goulart BH, Anderson LN, Manjarrez KL, Dietsch GN, Bryan JK, Hershberg RM, Disis ML, Martins RG. 2017. Phase Ib Trial of the Toll-like Receptor 8 Agonist, Motolimod (VTX-2337), Combined with Cetuximab in Patients with Recurrent or Metastatic SCCHN. Clin Cancer Res 23: 2442–50

35. Lu H, Dietsch GN, Matthews MA, Yang Y, Ghanekar S, Inokuma M, Suni M, Maino VC, Henderson KE, Howbert JJ, Disis ML, Hershberg RM. 2012. VTX-2337 is a novel TLR8 agonist that activates NK cells and augments ADCC. Clin Cancer Res 18: 499–509

36. Monk BJ, Brady MF, Aghajanian C, Lankes HA, Rizack T, Leach J, Fowler JM, Higgins R, Hanjani P, Morgan M, Edwards R, Bradley W, Kolevska T, Foukas P, Swisher EM, Anderson KS, Gottardo R, Bryan JK, Newkirk M, Manjarrez KL, Mannel RS, Hershberg RM, Coukos G. 2017. A phase 2, randomized, double-blind, placebo-controlled study of chemo-immunotherapy combination using motolimod with pegylated liposomal doxorubicin in recurrent or persistent ovarian cancer: a Gynecologic Oncology Group partners study. Ann Oncol 28: 996–1004

37. Ferris RL, Saba NF, Gitlitz BJ, Haddad R, Sukari A, Neupane P, Morris JC, Misiukiewicz K, Bauman JE, Fenton M, Jimeno A, Adkins DR, Schneider CJ, Sacco AG, Shirai K, Bowles DW, Gibson M, Nwizu T, Gottardo R, Manjarrez KL, Dietsch GN, Bryan JK, Hershberg RM, Cohen EEW. 2018. Effect of Adding Motolimod to Standard Combination Chemotherapy and Cetuximab Treatment of Patients With Squamous Cell Carcinoma of the Head and Neck: The Active8 Randomized Clinical Trial. JAMA Oncol 4: 1583–8

38. Stephenson RM, Lim CM, Matthews M, Dietsch G, Hershberg R, Ferris RL. 2013. TLR8 stimulation enhances cetuximab-mediated natural killer cell lysis of head and neck cancer cells and dendritic cell cross-priming of EGFR-specific CD8+ T cells. Cancer Immunol Immunother 62: 1347–57

39. Dietsch GN, Randall TD, Gottardo R, Northfelt DW, Ramanathan RK, Cohen PA, Manjarrez KL, Newkirk M, Bryan JK, Hershberg RM. 2015. Late-Stage Cancer Patients Remain Highly Responsive to Immune Activation by the Selective TLR8 Agonist Motolimod (VTX-2337). Clin Cancer Res 21: 5445–52

40. Northfelt DW, Ramanathan RK, Cohen PA, Von Hoff DD, Weiss GJ, Dietsch GN, Manjarrez KL, Randall TD, Hershberg RM. 2014. A phase I dose-finding study of the novel Toll-like receptor 8 agonist VTX-2337 in adult subjects with advanced solid tumors or lymphoma. Clin Cancer Res 20: 3683–91

41. Shayan G, Kansy BA, Gibson SP, Srivastava RM, Bryan JK, Bauman JE, Ohr J, Kim S, Duvvuri U, Clump DA, Heron DE, Johnson JT, Hershberg RM, Ferris RL. 2018. Phase Ib Study of Immune Biomarker Modulation with Neoadjuvant Cetuximab and TLR8 Stimulation in Head and Neck Cancer to Overcome Suppressive Myeloid Signals. Clin Cancer Res 24: 62–72

42. Cervantes JL, Weinerman B, Basole C, Salazar JC. 2012. TLR8: the forgotten relative revindicated. Cell Mol Immunol 9: 434–8

43. Leijen S, van Geel RM, Sonke GS, de Jong D, Rosenberg EH, Marchetti S, Pluim D, van Werkhoven E, Rose S, Lee MA, Freshwater T, Beijnen JH, Schellens JH. 2016. Phase II Study of WEE1 Inhibitor AZD1775 Plus Carboplatin in Patients With TP53-Mutated Ovarian Cancer Refractory or Resistant to First-Line Therapy Within 3 Months. J Clin Oncol 34: 4354–61

44. Greulich W, Wagner M, Gaidt MM, Stafford C, Cheng Y, Linder A, Carell T, Hornung V. 2019. TLR8 Is a Sensor of RNase T2 Degradation Products. Cell 179: 1264–75 e13

45. Brueseke TJ, Tewari KS. 2013. Toll-like receptor 8: augmentation of innate immunity in platinum resistant ovarian carcinoma. Clin Pharmacol 5: 13–9

46. Ye J, Ma C, Hsueh EC, Dou J, Mo W, Liu S, Han B, Huang Y, Zhang Y, Varvares MA, Hoft DF, Peng G. 2014. TLR8 signaling enhances tumor immunity by preventing tumor-induced T-cell senescence. EMBO Mol Med 6: 1294–311

47. Gorden KK, Qiu XX, Binsfeld CC, Vasilakos JP, Alkan SS. 2006. Cutting edge: activation of murine TLR8 by a combination of imidazoquinoline immune response modifiers and polyT oligodeoxynucleotides. J Immunol 177: 6584–7

48. Bracci L, Schiavoni G, Sistigu A, Belardelli F. 2014. Immune-based mechanisms of cytotoxic chemotherapy: implications for the design of novel and rationale-based combined treatments against cancer. Cell Death Differ 21: 15–25

49. Do K, Doroshow JH, Kummar S. 2013. Wee1 kinase as a target for cancer therapy. Cell Cycle 12: 3159–64

50. Kim H, Griffith TS, Panyam J. 2019. Poly(d,l-lactide-co-glycolide) Nanoparticles as Delivery Platforms for TLR7/8 Agonist-Based Cancer Vaccine. J Pharmacol Exp Ther 370: 715–24

